# Structural instability and divergence from conserved residues underlie intracellular retention of mammalian odorant receptors

**DOI:** 10.1101/605337

**Authors:** Kentaro Ikegami, Claire A. de March, Maira H. Nagai, Soumadwip Ghosh, Matthew Do, Ruchira Sharma, Elise S. Bruguera, Yueyang Eric Lu, Yosuke Fukutani, Nagarajan Vaidehi, Masafumi Yohda, Hiroaki Matsunami

## Abstract

Mammalian odorant receptors are a diverse and rapidly evolving set of G protein-coupled receptors expressed in olfactory cilia membranes. Most odorant receptors show little to no cell surface expression in non-olfactory cells due to endoplasmic reticulum retention, which has slowed down biochemical studies. Here, we provide evidence that structural instability and divergence from conserved residues of individual odorant receptors underlie intracellular retention using a combination of large-scale screening of odorant receptors cell surface expression in heterologous cells, point mutations, structural modeling, and machine learning techniques. We demonstrate the importance of conserved residues by synthesizing “consensus” odorant receptors that show high levels of cell surface expression similar to conventional G protein-coupled receptors. Furthermore, we associate *in silico* structural instability with poor cell surface expression using molecular dynamics simulations. We propose an enhanced evolutionary capacitance of olfactory sensory neurons that enable the functional expression of odorant receptors with cryptic mutations.

**Significance Statement:** Odor detection in mammals depends on the largest family of G protein-coupled receptors, the odorant receptors, which represent ∼2% of our protein-coding genes. The vast majority of odorant receptors are trapped within the cell when expressed in non-olfactory cells. The underlying causes of why odorant receptors cannot be functionally expressed in non-olfactory cells have remained enigmatic for over 20 years. Our study points to divergence from a consensus sequence as a key factor in a receptor’s inability to function in non-olfactory cells, which in turn, helps explain odorant receptors’ exceptional functional diversity and rapid evolution. We also show the success of protein engineering strategies for promoting odorant receptor cell surface expression.

---

Mammalian olfactory receptors (ORs) are the largest and most diverse family of G protein-coupled receptors (GPCRs) (1, 2). ORs are expressed on the cell surface of olfactory sensory neurons (OSNs) to detect and discriminate the vast number of odors in the environment (3, 4). ORs are rapidly evolving with gene duplications and deletions, in addition to functional modifications between homologs presumably for species-specific environmental adaptation (2, 5, 6). ORs are usually retained in the endoplasmic reticulum (ER) when expressed alone in non-olfactory cells, including neurons (7–14).

Receptor transporting protein (RTP) 1 and RTP2, which are proposed to act as chaperones in the olfactory sensory neurons, enhance the cell surface expression of many ORs when co-transfected in heterologous cells (15–20). In *RTP1* and *RTP2* double knockout mice (RTP DKO), the majority of ORs are significantly underrepresented (uORs) due to the absence of mature OSNs expressing them, suggesting that these ORs require RTP1 and RTP2 in order to function (21). Interestingly, a small subset of ORs is overrepresented (oORs), suggesting that a minor subset of ORs function without RTP1 and RTP2. Accordingly, some oORs show cell surface expression when expressed without RTPs in heterologous cells (21).

The underlying causes of OR retention in the ER in cells other than mature OSNs, as well as how RTP1 and RTP2 promote OR trafficking are not well understood. Adding export signals or making OR chimeras with canonical GPCRs have been shown to enhance functional expression of some ORs. However, previous structure-functional analysis using model ORs based on the assumption of OR-specific ER retention signals did not identify common residues that are involved in the cell surface expression of ORs (9, 22, 23).

In this study, we approach the mechanistic understanding of OR trafficking with the goals of identifying specific residues underlying ER retention and, using this knowledge, engineering ORs with increased expression in heterologous cells similar to that of non-olfactory GPCRs. To achieve these goals, we have used inter-disciplinary strategies. First, we used a pair of closely related ORs that show differential cell surface expression in heterologous cells to identify specific amino acid residues that influence cell surface expression. We performed molecular dynamics (MD) simulations on a set of ORs and mutants with differential cell surface expression to estimate protein stability and its possible relationship to expression. Second, we conducted a large-scale analysis of the cell surface expression of 210 ORs. We used the dataset to identify critical residues from which we built a machine-learning model to predict cell surface expression. Third, we synthesized ORs based on insights from the model to demonstrate the role of conserved residues in OR trafficking. Fourth, stabilization strategies commonly used on GPCRs and other proteins (24–27) were applied to ORs. We improved the stability of the most promising consensus ORs by inserting salt bridges in their structure and obtained mutated consensus ORs that show surface expression levels comparable to a canonical GPCR. Together, our data suggest that divergence from conserved residues results in the retention of ORs inside the cells, which may be caused by structural instability. We hypothesize that an enhanced evolutionary capacitance in the olfactory sensory neurons with olfactory-specific chaperones would enable rapid functional evolution of ORs (28–32).

## Results

### A TM4 residue, G^4.53^, is crucial for cell surface trafficking of model ORs

All OR cell surface expressions have been evaluated by flow cytometry (see Method section). We chose Olfr539 and Olfr541 with 90% amino acid identity as a model system to study OR trafficking. Olfr539 is an oOR, an indication of high representation in the absence of RTP1 and RTP2 *in vivo*, that shows robust cell surface expression in human embryonic kidney (HEK) 293T cells. In contrast, Olfr541 is an uOR, which requires RTP1 and RTP2 for its cell surface expression *in vivo*, that shows no detectable cell surface expression in HEK293T cells (Fig. 1A, 1B). We generated a series of chimeric ORs by intermingling parts of the amino acid sequence of Olfr539 with that of Olfr541 (Fig. 1C). Cell surface expression levels of the chimeric ORs were measured by flow cytometry. Mutant ORs which had the middle region (from 152^th^ residue to 247^th^ residue) of Olfr539 showed high surface expression levels, indicating an important domain for cell surface expression.

**Figure 1.**
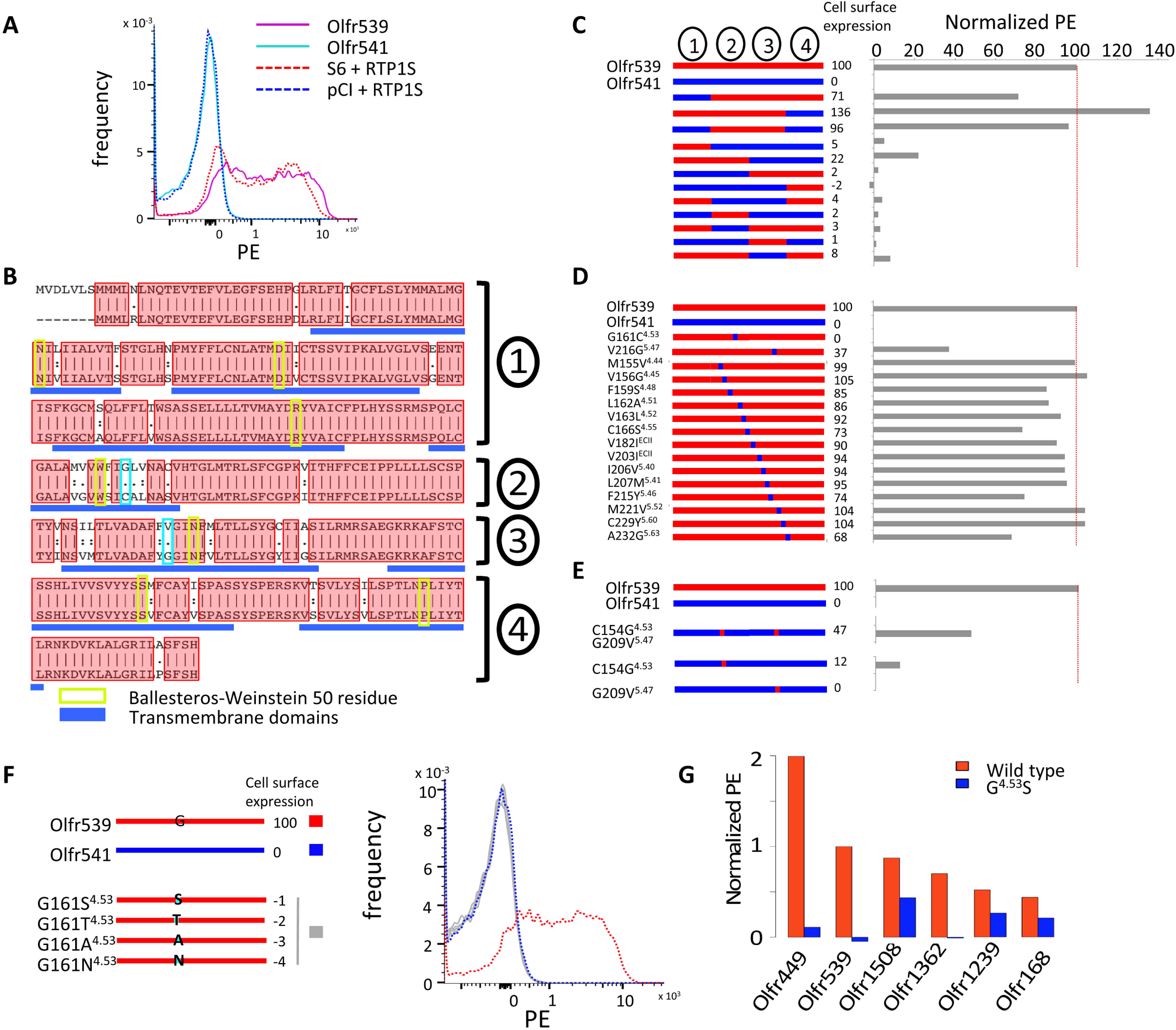
G161^4.53^ and V216^5.47^ are critical to the cell surface expression of Olfr539 and Olfr541. **A**, Olfr539 robustly but Olfr541 poorly expresses in the cell surface in heterologous cells in the absence of RTP1 and RTP2. The expression is evaluated by flowcytometry with OR S6 and pCI as positive and negative controls, respectively, by recording the frequency of PE fluorescence (see Methods section). **B**, Alignment of protein sequences of Olfr539 (top) and Olfr541 (bottom). 90% of the amino acid residues are shared between the two ORs and are shown in red boxes. Ballesteros-Weinstein 50 residue for each transmembrane domain is boxed in green (33). G/C161^4.53^ and G/V216^5.47^ are highlighted in cyan boxes. **C, D, E, F**, Designs of chimeric ORs are shown on the left and cell surface expression results (Normalized PE) on the right. **C**, Chimeric ORs created by replacing parts of Olfr539 (red) with those of Olfr541 (blue) **D**, Single amino acid mutants created by substituting amino acids of Olfr539 in region 2 and 3 for those of Olfr541. **E**, Reciprocal Olfr541 mutants created by substituting single or double amino acids of Olfr541 with those of Olfr539. **F**, Olfr539 mutants created by substituting G161^4.53^ with S, T, A or N, which are altogether conserved in almost 30% of the mouse ORs, lose cell surface expression. **G**, G^4.53^S mutants of RTP-independent ORs (Olfr449, Olfr539, Olfr1508, Olfr1362, Olfr1239 and Olfr168) (blue) show less cell surface expression levels compared with the wild types (red) in the absence of RTP1 and RTP2.

We next generated single amino acid mutants of each middle region residue of Olfr539 by substituting each amino acid of Olfr539 with that of Olfr541 (Fig. 1D). Olfr539 G161C^4.53^ (4.53 refers to the Ballesteros-Weinstein residue numbering system used for GPCRs (33), see Fig. 1B) and V216G^5.47^ showed abolished or diminished Olfr539 cell surface expression. Accordingly, we expressed the reciprocal single mutants Olfr541 C154G^4.53^ and Olfr541 G209V^5.47^, and the double mutant Olfr541 C154G^4.53^/G209V^5.47^ to test if these residues are sufficient for promoting cell surface expression. Olfr541 C154G^4.53^ showed a moderate level of cell surface expression, although Olfr541 G209V^5.47^ did not show any improvements (Fig. 1E). Olfr541 C154G^4.53^/G209V^5.47^ showed a more enhanced cell surface expression than C154G^4.53^ alone, suggesting a synergistic interaction between the two critical residues in facilitating cell surface trafficking. This indicated that these two residues are critical in determining cell surface expression levels. Other residues had little or no effect on cell surface expression.

We further investigated which properties of G^4.53^ were influencing with controlling the cell surface trafficking. We generated mutants by substituting G161^4.53^ of Olfr539 for less frequently used amino acids (S, T, A or N), which were altogether present in almost 30% of mouse ORs (Fig. 1F). None of these mutants exhibited cell surface expression, indicating a requirement of G^4.53^ for the cell surface trafficking of Olfr539. We next tested whether G^4.53^ controlled the trafficking of other ORs that are trafficked to the cell surface. We chose five ORs (Olfr1362, Olfr1508, Olfr449, Olfr168 and Olfr1239) that show cell surface expression in HEK293T cells (RTP-independent ORs, see below for details). All the G^4.53^S mutants showed abolished or diminished cell surface expression (Fig. 1G) supporting the importance of this residue in facilitating cell surface trafficking of ORs.

### Contribution of G^4.53^ in the *in silico* structural stability of Olfr539 and Olfr541

In order to understand how the two critical residues, G^4.53^ and V^5.47^, influence the cell surface trafficking of Olfr539 and Olfr541, we localized the two residues in 3D structural models of Olfr539 and Olfr541. OR models were constructed using a homology modeling method based on known GPCR experimental structures in the inactive state (34, 35). G^4.53^ and V^5.47^ are in the middle of the 4^th^ and the 5^th^ transmembrane domains (TM), respectively (Fig. 2A). Following the structural model, G^4.53^ is unlikely to directly interact with odorants as TM4 is not involved in the binding cavity. However, V^5.47^ is part of TM5 and located at the cradle of the binding site. To evaluate if our mutations change the selectivity of Olfr539, we performed an activation screening on Olfr539wt and G4.53 (C,S,T,A and N) and V5.47 (G) mutants against 22 odorants (Fig. S1). We saw that, if the receptor responds to any of the tested odorants, the response profiles of the mutated receptors are similar to those of the wt Olfr539, suggesting that the selectivity of the receptor is not grossly changed and the residues 4.53 and 5.47 were not directly involved in odorant binding.

**Figure 2.**
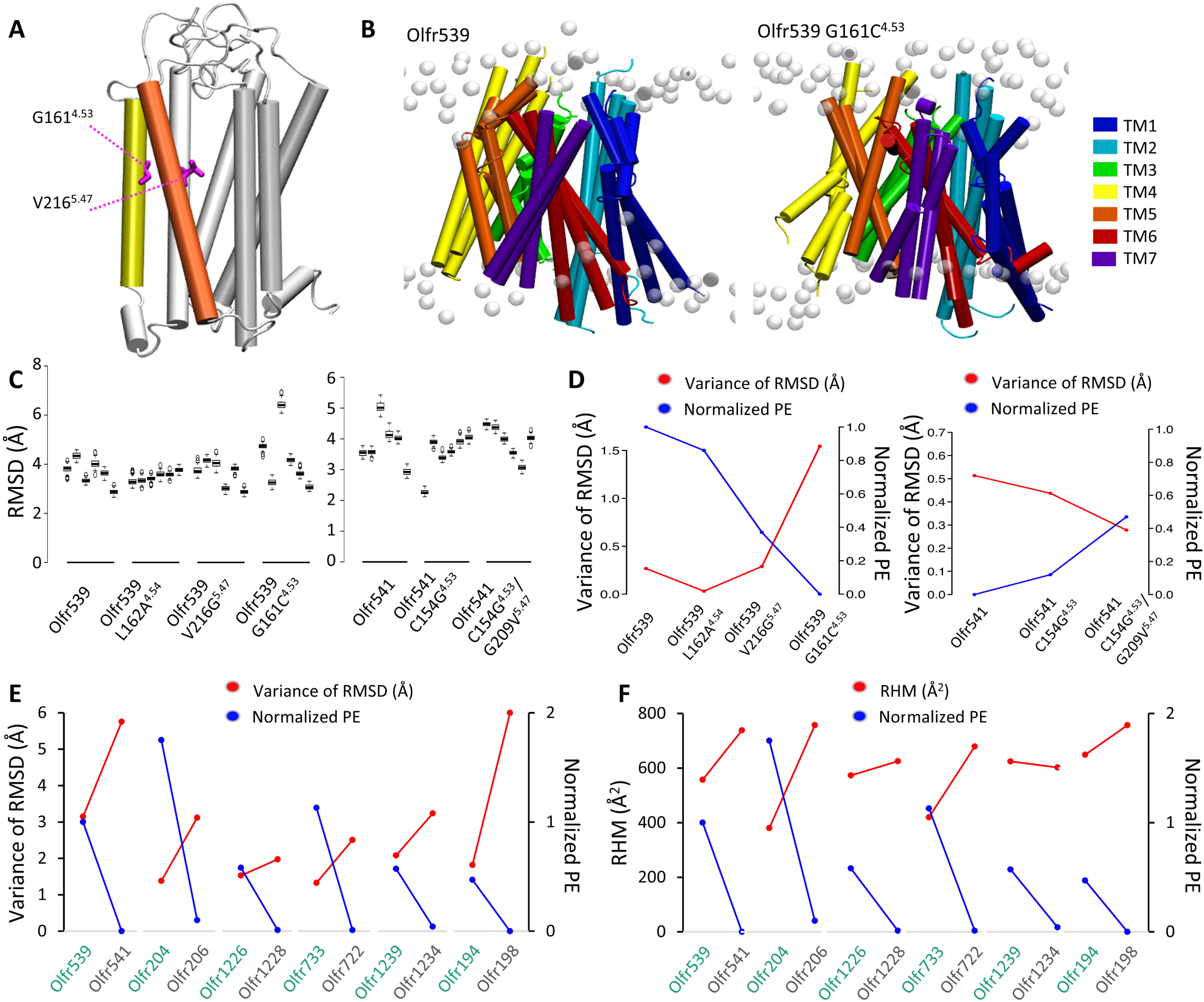
*In silico* structural stability of ORs correlates with the cell surface expression level **A**, 3D homology model of Olfr539. TM4 (yellow) and TM5 (orange) are represented in colored tubes and the remaining structure is in white. G154^4.53^ and V209^5.47^ are developed in pink licorice. **B**, Superposed images of triplicate wild type Olfr539 (left) and Olfr539 G161C^4.53^ (right) models after 500ns molecular dynamics (MD) simulations in an explicit model of plasma membrane (white). **C**, RMSDs of 6 individual MD simulations of Olfr539 systems (wild type, L162A^4.54^ G161C^4.53^ and V216G^5.47^) (left), and Olfr541 systems (wild type, C154G^4.53^ and C154G^4.53^/G209V^5.47^). The models are placed in descending order based on cell surface expression levels for Olfr539 systems and in ascending order for Olfr541 systems. **D**, Plots of the variance of mean RMSDs (left axis, red plots) and the cell surface expression levels (right axis, blue plots) of Olfr539 systems (left) and Olfr541 systems (right). **E**, Plots of the variance of RMSDs (left axis, red plots) and the cell surface expression levels (right axis, blue plots) of Olfr pairs sharing a high sequence identity but showing different cell surface expression levels (high in green, low in gray). **F**, Plots of the Residual Hydrophobic Mismatch (RHM, left axis, blue plots) and the cell surface expression levels (right axis, blue plots) of Olfr pairs sharing a high sequence identity but showing different cell surface expression levels (high in green, low in gray).

We hypothesized that diminished structural stability of ORs might cause ER retention due to general quality control mechanisms and that G^4.53^ and V^5.47^ are important for the stabilization of ORs. To test this, we built Olfr539, Olfr541, Olfr539 G161C^4.53^, Olfr539 V216G^5.47^, Olfr541 C154G^4.53^, and Olfr541 C154G^4.53^/G209V^5.47^ homology models. We also included a control mutant Olfr539 L162A^4.54^ which shows a similar cell surface expression to Olfr539. We ran six independent MD simulations of 500ns for all the systems embedded in an explicit lipid bilayer. All six simulations of Olfr539 showed well-packed structures while Olfr539 G161C^4.53^ showed flexibility (Fig. 2B and C) as quantified by the root mean square deviations (RMSD) of atomic positions of TM domains for each MD simulation. Olfr539 and L162A^4.54^ showed similar RMSD across multiple simulations, all converging to equivalent equilibrated structures. However, Olfr539G161C^4.53^ and V216G^5.47^, which are poorly expressed, showed variability in structures. This suggests that poorly trafficked ORs have more flexible structures and are unstable when inserted into a cell membrane model (Fig. 2C, S2). In contrast, Olfr541 showed a wide range of RMSD, whereas its mutants C154G^4.53^ and C154G/G209V showed less variation in RMSD (Fig. 2C). To further examine whether structural stability is associated with cell surface expression level, we plotted the cell surface expression levels and variances of RMSDs for both Olfr539 and Olfr541 systems. Indeed, the mutants with lower cell surface expression showed a larger variance of RMSDs. Conversely, the mutants with higher cell surface expression showed a smaller variance of RMSDs (Fig. 2D). This trend is more pronounced in RMSDs calculated for the extracellular (EC) side of the TM domains, suggesting that stabilities of this part may be especially important for cell surface expression (Fig. S3). We also plotted the cell surface expression levels and variances of RSMD for five other OR pairs that show a high percentage of sequence identity but different cell surface expression levels (Fig. 2E, Fig S3). We found again a clear anti-correlation between the variance of RMSD and cell surface expression. To evaluate further the structural correlation with cell surface expression between these pairs, we calculated the Residual Hydrophobic Mismatch (RHM) between the OR and the membrane during our MD simulations (Fig. 2F). We found that five out of six OR pairs show an anti-correlation between RHM and the cell surface expression. Further, we identified a linear correlation between differences of RHM and differences of cell surface expression in each pair (Fig. S3, R^2^=0.9035). All together, these data support the hypothesis that structural stability contributes to the cell surface expression of ORs.

### A comprehensive evaluation of cell surface expression levels of ORs in heterologous cells

So far, we showed that the TM4 residue at the position 4.53 plays a crucial role in OR trafficking. However, it is unlikely that this position is the sole determinant, as G^4.53^ is present in 66% of mouse ORs, yet the vast majority of ORs do not show cell surface expression in heterologous cells. To gain a more comprehensive understanding of OR cell surface expression, we selected 210 ORs (76 oORs and 134 uORs *in vivo*) and tested which ORs exhibit cell surface expression in heterologous cells. Consistent with previous findings (21), oORs, as a group, showed more robust OR surface expression than uORs (p<0.05, U test) (Fig. 3A). Using the approximation that surface expression levels are the overlap of two normal distributions of positive ORs and negative ORs, we defined the ORs with expression levels of more than 0.144 (A.U.) as positive ORs (cut off = top 0.001 of negative ORs). 34/210 ORs (14.0%) were positive under this criterion. Consistent with the more robust cell surface expression of oORs, 26/76 (34.2%) of oORs and only 8/134 (6.0%) of uORs showed positive cell surface expression. We defined the 26 ORs that are oORs and cell surface expression positive as <u>RTP-independent</u> ORs and the 126 ORs that are uORs and cell surface expression negative as <u>RTP-dependent</u> ORs.

**Figure 3.**
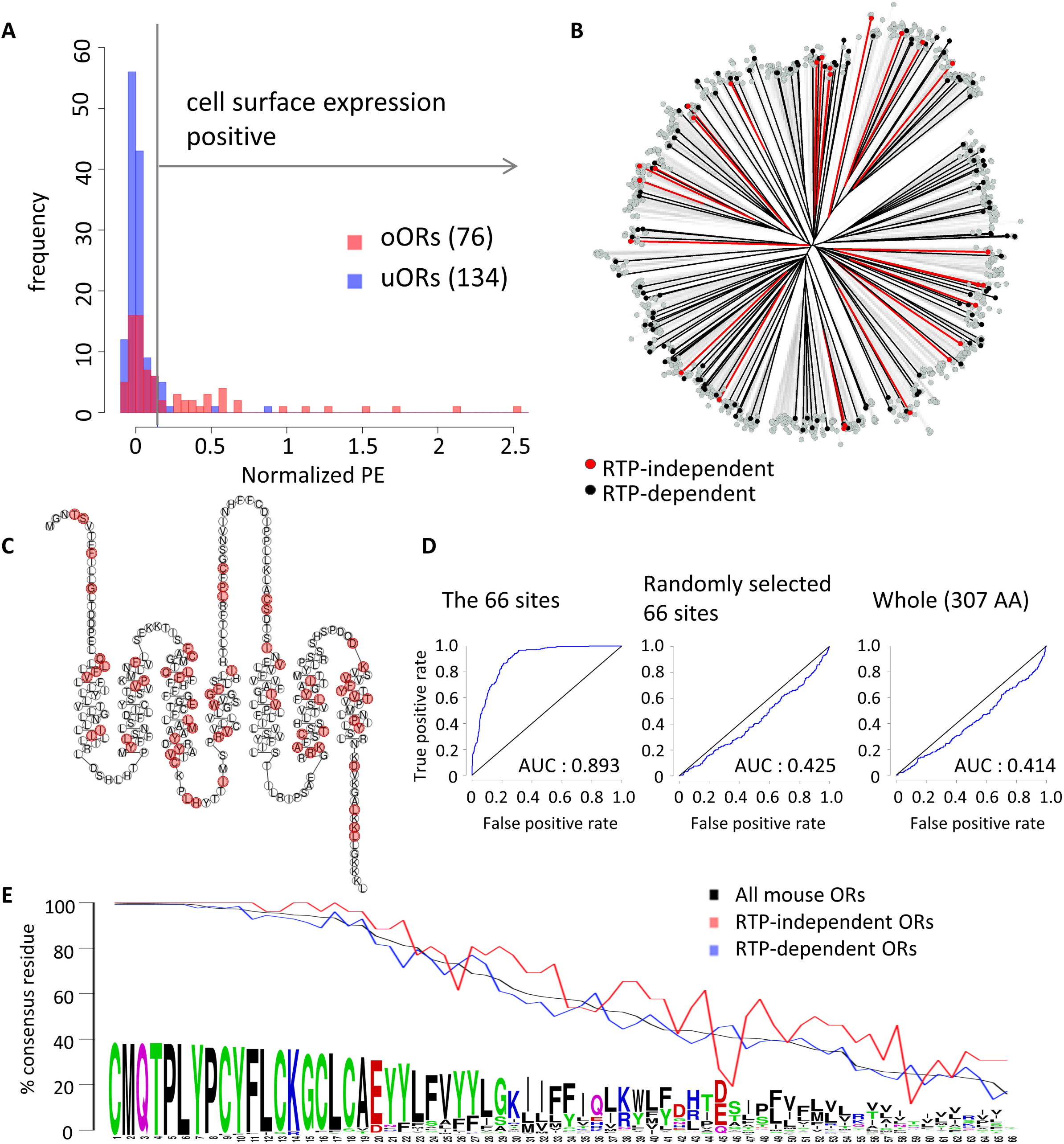
Cell surface expression analyses for a large repertoire of mouse ORs in heterologous cells in the absence of RTP1 and RTP2. **A**, Histogram of cell surface expression levels of 76 oORs (red) and 134 uORs (blue). 0.144 (vertical line) is a cut-off for positive/negative cell surface expression. **B**, Phylogenetic tree of protein sequences of mouse ORs. Red, black and gray indicate RTP-independent ORs, RTP-dependent ORs and the others ORs respectively. **C**, Snake plot of the consensus protein sequence of mouse ORs. The 66 sites with less diverse residues in RTP-independent ORs (p<0.05, Bonferroni corrected) are colored in red. **D**, RTP dependence of tested ORs is predicted using SVM-based classifiers based on amino acid properties in ten-fold cross-validation, and the accuracy is validated by the ROC curves. The SVM-based classifier has been built using the 66 sites of interest, 66 random amino acid positions and the entire 307 amino acid positions in the tested ORs. **E**, Usage rate of consensus residues at the 66 sites. RTP-independent ORs more frequently use consensus residues than RTP-dependent ORs at 58 out of 66 sites.

### Critical residues predict cell surface expression of ORs

We hypothesized that residues specific to RTP-independent or RTP-dependent ORs are critical for OR trafficking in the absence of RTP1 and RTP2. To investigate whether overall amino acid sequence similarities are associated with cell surface trafficking, we asked whether RTP-independent and/or RTP-dependent ORs are clustered in a phylogenetic tree (Fig. 3B, S4). Both are distributed on multiple branches, indicating that overall similarities do not determine their ability to be trafficked to the cell surface.

Next, we hypothesized that amino acid residues at specific positions create a network that controls cell surface trafficking. To investigate this idea, we aligned 26 RTP independent and 126 RTP dependent ORs (total of 152 ORs) and calculated Grantham distances (36) of amino acid properties at individual sites. We identified 66 positions with lower Grantham distances between amino acids for RTP-independent ORs than for all the 152 tested ORs (cut off p<0.05, t-test with Bonferroni correction). As expected, the position 4.53 is one of these 66 sites; 80.8% of RTP-independent ORs possess a G residue at this position against only 61.1% in the RTP-dependent ORs.

Contrary to the initial assumption that specific domains control OR cell surface expression, the 66 sites were scattered throughout the OR sequence. Moreover, there was no specific site that was exclusively present in one of the groups, suggesting that there are no trafficking promotion or inhibition signals that are shared among all ORs (Fig. 3C). To investigate whether these 66 sites can predict the RTP dependence of ORs, we classified tested ORs by support-vector machine-based classifiers in ten-fold cross-validation. The support vector machine model generated by the 66 sites of amino acid residues discriminated RTP-independent ORs (1.70 x 10^-92^, Wilcoxon signed rank test; AUC = 0.893). However, those generated by the 66 randomly selected sites (p=0.999, Wilcoxon signed rank test; AUC = 0.425) and those generated by all sites (p=0.999, Wilcoxon signed rank test; AUC = 0.414) failed to discriminate RTP-independent ORs. This demonstrates that these 66 sites robustly predict whether an OR shows cell surface expression in heterologous cells (Fig. 3D).

What properties of these 66 residues are associated with cell surface expression? When we looked at the degree of conservation of these residues, many of these sites were conserved among ORs (Fig. 3E, S5). 29% of the sites (19 / 66) are conserved in more than 90% of the mouse ORs whereas only 13% of all amino acid sites (41/307) are conserved in more than 90% of full-length ORs (p=0.0048, Fisher’s exact test). RTP-independent ORs have the most common amino acid residues much more frequently present than RTP-dependent ORs (58 out of the 66 sites, p=6.35×10^-6^, chi-square test), suggesting that ORs that are in line with consensus amino acids in these positions are more likely to show cell surface expression.

### Engineered consensus ORs robustly express on the cell surface in heterologous cells

The above results suggest the importance of the most frequently occurring amino acid at a given site in cell surface expression. This observation led us to predict that ORs that are designed based on “consensus” amino acids for each site would be efficiently trafficked to the cell surface. The consensus strategy has already been applied to proteins or codons to improve their thermostability or function in other proteins (24, 25, 37, 38). The success of this strategy relies on the number of proteins available to build the consensus sequence and their sequence similarity (24, 25). Here, we used the unique diversity of the OR family among GPCRs to apply the consensus strategy, aiming to obtain stable ORs. We aligned amino acid sequences of human OR families and determined the consensus sequences as the most frequently occurring amino acid residue at each position (Table S1). We first chose the OR10 family to measure the cell surface expression levels of the consensus OR in HEK293T cells in comparison to each member of OR10 family. Strikingly, the consensus OR10 robustly expressed on the cell surface, more than any of the individual OR10 family members tested (Fig. 4A). We generated consensus human ORs for 8 other families (OR1, OR2, OR4, OR5, OR6, OR8, OR51 and OR52) and measured their cell surface expression levels in the absence of RTPs. 6 out of 9 consensus ORs show more robust cell surface expression than any tested natural ORs (Fig. 4B).

**Figure 4.**
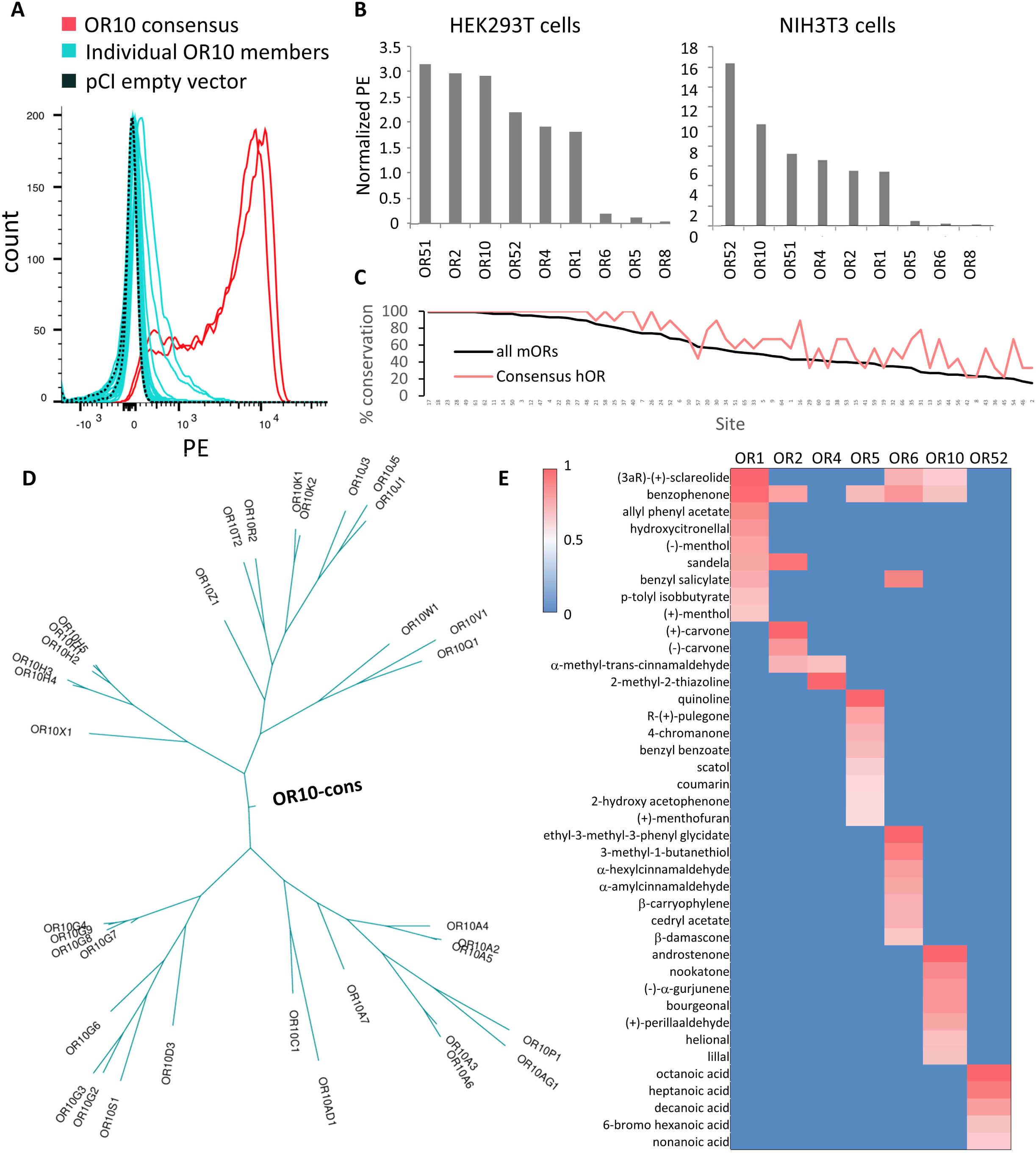
Potential of consensus ORs for robust cell surface expression in the absence of RTP1S. **A**, 32 human OR10 subfamily members (cyan) and the consensus OR (red, duplicate) were transfected into HEK293T cells and their cell surface expression levels were measured by flowcytometry. **B**, Cell surface expression levels of 9 consensus ORs (OR1, OR2, OR4, OR5, OR6, OR8, OR10, OR51 and OR52) were evaluated in HEK293T or NIH/3T3 cells. The PE fluorescence is normalized by setting Olfr539 response to 1 and Olfr541 to 0. **C**, Usage rate of consensus residues at the 66 sites for consensus human ORs (Consensus hOR, red) and all mouse ORs (black). **D**, Ancestral tree of OR10 family member including OR10-consensus. **E**, Heatmap of ORs’ responses to 50 µM of odorants selected from a previously screened panel of 320 compounds. Luciferase activity was normalized for each OR by setting as 1.0 the highest response value, and 0 as the lowest response value. Consensus OR1, OR2, OR4, OR5, OR6, OR10 and OR52 are functionally expressed in Hana3A cells.

To exclude the possibility that this effect was specific only to HEK293T cells, we expressed the consensus ORs in NIH3T3 cells, which are derived from mouse fibroblasts. Again, most of the consensus ORs showed robust cell surface expression (Fig. 4B). We evaluated the level of conservation of the 66 amino acid sites that we showed to be important for RTP-independent OR expression between our consensus ORs and the mouse OR repertoire (Fig. 3E). Consensus ORs have the most common amino acid residues much more frequently represented than natural ORs (59 out of the 66 sites, p=1.24×10^-9^, Wilcoxon signed rank test) (Fig. 4C). We built phylogenetic trees using parsimony criterion for each OR family including the corresponding consensus OR and found that the consensus OR is always located at the origin of the tree (Fig. 4D, S6).

Finally, we verified that the consensus receptors are functional in their response to odorants(19). We used cAMP-mediated luciferase reporter gene assay (16) to screen active ligands from a set of 320 diverse odorants at 50μM. We identified robust ligands for OR1, OR2, OR4, OR5, OR6, OR10 and OR52, each of which shows responses to specific subsets of the tested odorants (Fig. 4E). Our data shows that these consensus ORs are indeed functional suggesting a proper folding.

### Stabilization of consensus ORs structure by introduction of salt bridges

The consensus ORs already show a clear increase in cell surface expression in comparison to naturally occurring ORs. Since we observed that the expression of an OR seems to be correlated with its stability and rigidity when inserted in a membrane using MD simulations (Fig. 2B, C and D), we attempted to further improve the expression level of the consensus ORs by engineering salt bridges in the structures guided by 3D homology models (Fig. 5A). Thermostabilizing studies on β1-adrenergic receptor (39) and chemokine receptor (26) showed that tightening of interactions between intracellular loop 1 (ICL1) and the helix 8 improved the GPCR stabilization. Another study (40) showed that enriching the number of basic residues in the ICL of an OR enhances its expression. Based on these premises, we inserted triple Arg mutations in ICL1 and a negatively charged residue in helix 8 to promote salt bridge interactions between ICL1 and helix 8 in four human consensus ORs, namely OR1, OR10, OR51 and OR52 (Fig. 5B). We evaluated the cell surface expression of the consensus ORs with their corresponding mutants, including a well-studied non-olfactory class A GPCR, the muscarinic 3 receptor (M_3_), as a positive control for high expression (Fig. 5C). We observed enhanced expression of OR10 and OR52 mutants which are comparable to the expression level of the M_3_ receptor, even when we decreased the amount of DNA used in transfection by 100-fold. However, we did not observe any enhancement in the expression level of OR1 and OR51 mutants. The insertion of such stabilizing interactions might improve the rigidity of the OR structure that seems to aid expression. We tested their functionality in comparison to their corresponding consensus OR (Fig. 5D). Mut-OR10 and Mut-OR52 both responded to their agonists (androstenone and nonanoic acid respectively). Mut-OR52 showed activity similar to the consensus OR52, but the response of Mut-OR10 was diminished compared to OR10. We optimized the luciferase assay protocol to these unusually highly expressed ORs by testing different DNA concentrations for cell transfection. For both OR10/Mut-OR10 and OR52/Mut-OR52, decreasing the amount of transfected DNA by 10 to 100 fold compared with the optimized amount for natural ORs resulted in robust responses against tested odorants, showing the capacity of these consensus ORs in supporting high levels of functional expression.

**Figure 5.**
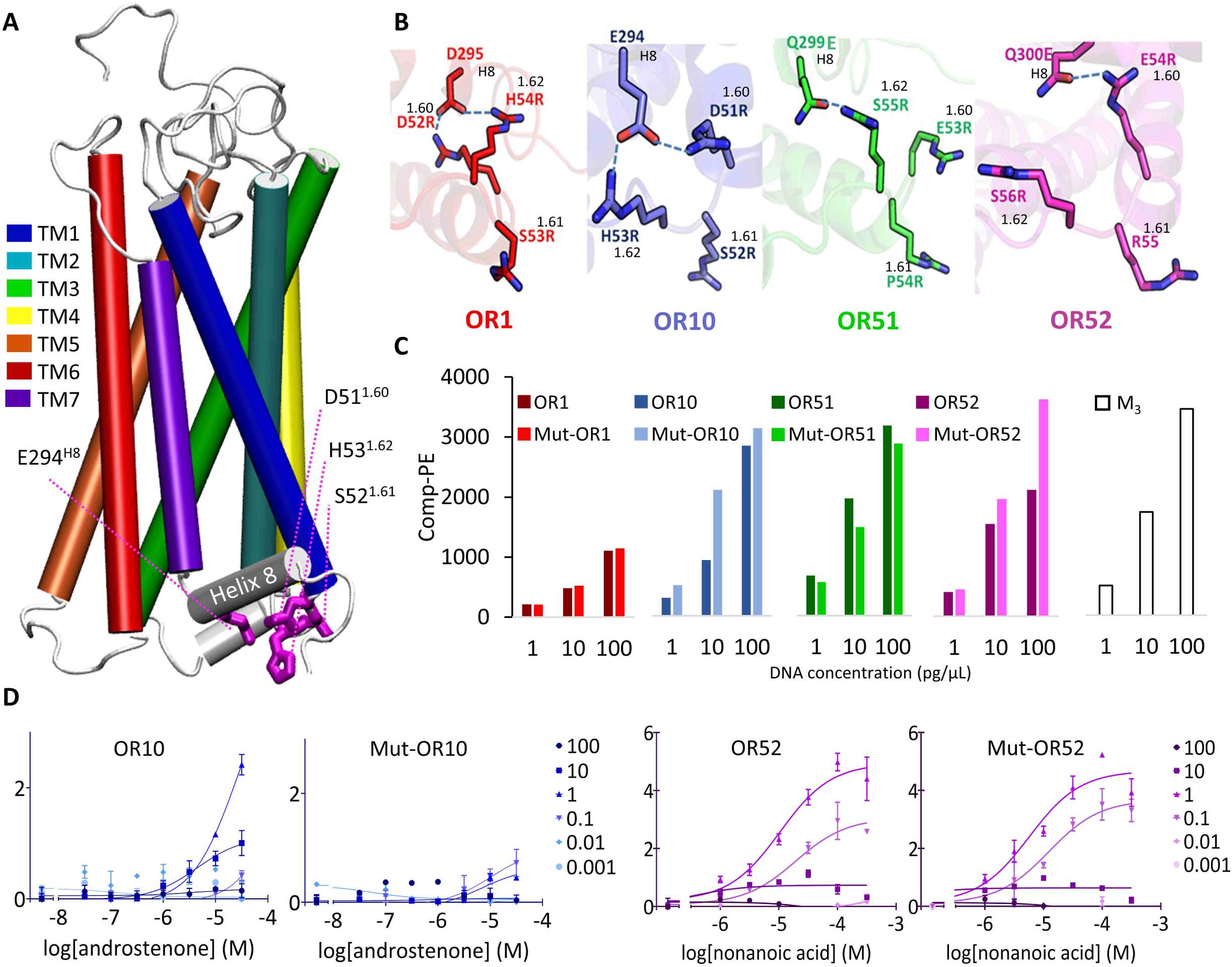
Improvement of OR expression by mutations in TM1 and helix 8. **A**, Homology model of OR10. Each TM is highlighted in a colored tube and residues D51^1.60^, S52^1.61^, H53^1.62^ and E294^H8^ are represented in licorice (pink). **B**, Zoom on the residues 1.60, 1.61, 1.62, H8 mutated for OR1 (red), OR10 (blue), OR51 (green) and OR52 (pink). Ionic interactions between the residues are shown by dotted lines. **C**, Expression analysis of OR1, OR10, OR51, OR52 and their mutants and the muscarinic receptor 3 (M3) at 1, 10 and 100 pg/µ L of DNA in the transfection mix. **D**, Dose-response curve of OR10 and OR52 and their mutants at different DNA concentration (from 0.001 to 100 in pg/µL of transfection mix) to androstenone and nonanoic acid, respectively. The y-axis represents the luciferase luminescence normalized to the basal activity of each DNA concentration.

## Discussion

The mammalian OR family is a unique protein family with its large size, rapid evolution and poor expression in heterologous cells, which makes their function notoriously difficult to study. Here, we investigated the underlying mechanisms by which OR trafficking is regulated in heterologous cells. Using chimeras and point mutations with a pair of highly similar ORs, we identified critical residues that regulate cell surface expression. MD simulations suggest that these residues may affect the flexibility and stability of ORs. We also conducted a large-scale cell-based screening to comprehensively identify ORs that are expressed on the cell surface, leading to the identification of specific residues associated with their cell surface expression. Cell-surface-expression-positive ORs tend to have conserved residues at the critical sites. We investigated if consensus ORs express on the cell surface in heterologous cells and demonstrated that most of the consensus ORs show robust cell surface expression.

Which residues or domains make OR trafficking difficult? Previous studies suggest different residues, domains, or features of ORs are responsible for intracellular retention (9, 23). Among them, a study proposed that fewer charged residues and more hydrophobic residues distributed throughout ORs might underlie intracellular retention (22). We tested whether RTP-independent ORs possess fewer charged and more hydrophobic residues compared to RTP-dependent ORs, as suggested by Bubnell et al (22). We found no significant differences between tested ORs (p=0.74 and p=0.71, respectively), suggesting that these features do not explain the cell surface trafficking of ORs as a group. Our current study identified two sites (G^4.53^ and V^5.47^) contributing to cell surface expression in model ORs. In non-olfactory class A GPCRs, position 4.53 is conserved as S39%, A33%, V8%, T4%, C3.5%, P3%, G3%, I2%, M1.5% and position 5.47 is conserved as F65%, Y11%, L10%, N3.5%, V3%, G2%, I2%, C2%. Position 4.53, as well as position 2.47, has been identified as a conserved packing cluster center in class A GPCRs and its mutation in leucine (A^4.53^L) in Rhodopsin disrupts the structure of the receptor (41). In the β2-adrenergic receptor, S^4.53^ belong to the motif S^4.53^xxxS^4.57^ which doesn’t participate to the functionality of the receptor and seems to be more involved in helix packing, maintaining the stability of the receptor (42). The position 5.47 appears to play a role in cannabinoid receptor (CB1R) stabilization as the crystal structure reported (PDB:5TGZ) containing T210^3.46^A+E273^5.37^K+T283^5.47^V+R340^6.32^E mutations shows enhanced protein homogeneity and thermostability (43). However, these residues were not identified in previous studies on ORs, it seems that different residues regulate the trafficking of different ORs. This raised a question of whether there are any common mechanisms that regulate OR trafficking. Our statistical analysis based on a large-scale cell surface expression analysis of hundreds of ORs identified 66 residues scattered throughout the receptor that play a critical role in OR cell surface expression. Although there was no single residue or domain that solely determined cell surface expression, we succeeded in building a machine learning model that reliably predicts OR cell surface expression based on these residues. This suggests that these 66 residues differentially contribute to trafficking efficiency of individual ORs.

What features of ORs cause their ER retention when expressed in non-olfactory cells? It was long hypothesized that ORs may possess specific and conserved ER retention signals (8, 9, 15, 22). However, a previous report shows that mutating the most highly conserved OR-specific amino acids into one of a non-olfactory GPCR that shows high cell surface expression did not enhance the trafficking of a model OR (22). Our data is also inconsistent with the idea that conserved OR-specific residues or motifs cause ER retention. In contrast, our study demonstrated that consensus ORs show robust cell surface expression, suggesting that such ER retention signals, if any, would not be shared by OR members. How do ORs (aside from a minor subset) possess the common feature of being retained inside the cells? Our study supports the model that the intracellular retention of ORs is caused by the structural instability of ORs, which is caused by divergence from conserved residues at the critical sites. In other words, the majority of ORs do not fold correctly in heterologous cells due to a divergence from conserved residues, thus they are trapped by the general protein quality control in the ER and cannot be trafficked to the plasma membrane. This model explains why many consensus ORs exhibit robust cell surface expression, while the vast majority of natural ORs do not show detectable cell surface expression in heterologous cells (44).

Why are ORs functional in OSNs but not in heterologous cells when expressed alone? Our study supports the idea that accessory proteins or chaperones expressed in the OSNs assist the folding of structurally unstable ORs (Fig. 6). In addition to RTP1 and RTP2, a previous study showed that an Hsp70 homolog enhanced the cell surface expression of an OR, suggesting the importance of chaperones for the cell surface expression of ORs (45). A possibility is that the unstable nature of ORs may be integral to proper OSN development (21, 46). Initiation of OR expression induces unfolded protein response (UPR) in developing OSNs, suggesting inefficient folding and ER accumulation of ORs in these cells (46, 47). RTP1 and RTP2 are induced by UPR signaling in developing OSNs, presumably allowing OR proteins to exit the ER and downregulate UPR (21, 46, 47). There is also the possibility that OSNs lack specific quality control proteins that are common in other cell types, allowing less stable ORs to be trafficked to the cell membrane where they are finally functional. This may resemble the case of the cystic fibrosis transmembrane conductance regulator (CFTR) where a single amino acid deletion (ΔF508 CFTR) causes its ER retention to be regulated by, among others, calreticulin or the Hsp90 co-chaperone Aha1. The downregulation of these chaperones results in the functional plasma membrane expression of ΔF508 CFTR (48). Outside of the olfactory sensory neurons, ORs are found to be ectopically expressed in tissues as diverse as heart, gut, and testis (49, 50). These ectopic ORs are unlikely to need RTPs to be functionally expressed in many of these non-olfactory tissues where no or very low RTP1 and RTP2 expressions are detected. We looked at the nature of the 66 positions in a set of four mouse and nineteen human ORs detected ectopically (Fig. S7)(49, 50). The 66 positions are not significantly more conserved in ectopic ORs than all mouse ORs (38 out of the 66 sites, p=0.398, Wilcoxon signed rank test). However, these results need to be taken with caution as the functional role of most of the ectopic ORs remains to be determined and their functional expression might depend on other types of chaperone yet to be discovered.

**Figure 6.**
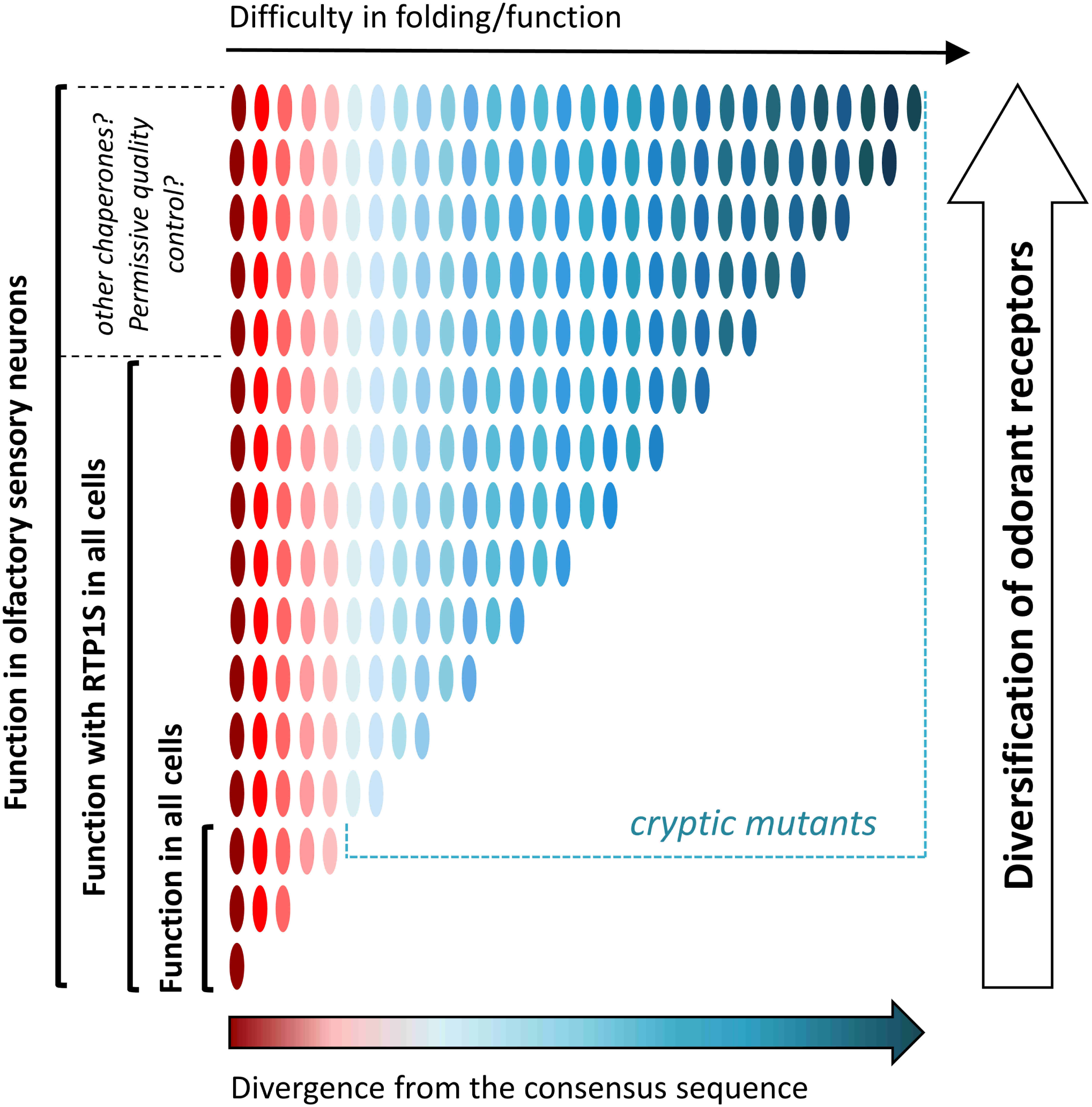
Proposed model. RTPs are evolutionary capacitors involved in enhanced diversification of ORs. OR genes are represented in oval shapes colored from dark red (ORs that are more aligned with the consensus) to dark blue (ORs that are more divergent from the consensus). Divergence of ORs from the consensus sequence results in difficulties in OR folding and function. Cryptic OR mutants are functional in the olfactory sensory neurons with RTPs and other capacitors. OR diversification and rapid evolution rely on the presence of olfactory-specific evolutionary capacitors.

Our study parallels with those conducted by Lindquist and others in deciphering the role of Hsp90, a chaperone that helps fold many proteins, as an evolutionary capacitor (Fig. 6) (29, 51, 52). Evolutionary capacitors, proteins that suppress deteriorating phenotypic variation under normal conditions, facilitate adaptation during evolution (31, 53). Our study suggests that RTP1, RTP2 and other chaperones yet to be discovered support functional expression of ORs that do not fold correctly in other cell types, conferring a unique capacity of the OSN. We speculate that olfactory-specific evolutionary capacitors play a role in the rapid evolution of ORs to facilitate sensory adaptation to detect and discriminate a vast number of odorants and natural odor mixtures (Fig. 6). Our model also implies that ORs in the ancestral species before expansion may resemble the consensus ORs thus they may not show similar levels of difficulties in cell surface expression in non-olfactory cells.

Lastly, despite some successes (10, 11), biochemical and structural studies of ORs have been challenging with the currently available expression systems, due to the relatively poor expression of ORs when compared to canonical GPCRs. Our study shows the success of the application of protein engineering strategies to enhance OR stability and cell surface expression. We have identified a set of engineered ORs that show robust cell surface expression comparable to the M_3_ muscarinic acetylcholine receptor, a canonical GPCR. The consensus ORs can serve as candidate ORs for future biochemical studies such as the large-scale production of ORs and purification. In addition, the residues regulating trafficking identified in this study will point to further improvements in protein production, which will accelerate our effort towards determining OR structures. Altogether, our study provides novel insights into how various residues in ORs regulate the receptor expression on the cell membrane.

## Methods

### DNA and vector preparation

Open reading frames of OR genes were subcloned into pCI (Promega) with a Rho tag at the N terminal. DNA fragments of OR genes were amplified by Phusion polymerase (Thermo Fisher Scientific). To generate chimeras and mutants of ORs, DNA fragments of OR genes were amplified by Phusion polymerase (Thermo Fisher Scientific). The fragments were mixed and amplified by PCR reaction to obtain full sequences. All plasmid sequences were verified using Sanger sequencing (3100 Genetic Analyzer, Applied Biosystems).

### Cell culture

HEK293T and Hana 3A cells (15) were grown in Minimal Essential Medium (MEM) containing 10% FBS (vol/vol) with penicillin-streptomycin and amphotericin B. Hana 3A cells were authenticated using polymorphic short tandem repeat (STR) at the Duke DNA Analysis Facility using GenePrint 10 (Promega) and shown to share profiles with the reference (ATCC). NIH/3T3 cells were grown in high glucose Dulbecco’s Modified Eagle Medium (DMEM) supplemented with 10% CS, penicillin-streptomycin and amphotericin B. All cell lines were incubated at 37°C, saturating humidity and 5% CO2. No mycoplasma infection was detected in all cell cultures.

### Flowcytometry analyses

The principle of the method can be found in Fig. S8. HEK293T cells were grown to confluency, resuspended and seeded onto 35 mm plates at 25% confluency. The cells were cultured overnight. A Rho tagged OR in the plasmid pCI and GFP expression vector were transfected using Lipofectamine 2000 (Fig. S8A and B). After 18-24 hours, the cells were resuspended by cell stripper and then kept in 5 mL round bottom polystyrene (PS) tubes (Falcon 2052) on ice. The cells were spun down at 4°C and resuspended in PBS containing 15 mM NaN3, and 2% FBS to wash the cell stripper. They were incubated in primary antibody (mouse anti Rho4D2 (54)) (Fig. S8C) and then washed, stained with phycoerythrin (PE)-conjugated donkey anti-mouse F(ab’)₂ Fragment antibody (Jackson Immunologicals: 715-116-150) (Fig. S8D) in the dark. To stain dead cells, 7-Amino-actinomycin D (Calbiochem) was added (Fig. S8E). The cells were analyzed using BD FACSCanto II FACS with gating allowing for GFP positive, single, spherical, viable cells, (Fig. S8F) and the measured PE fluorescence intensities were analyzed and visualized using Flowjo (55). We normalized the surface expression levels by cells expressing Olfr539, which was robustly expressed on the cell surface, and cells expressing Olfr541, which showed no detectable cell surface expression.

### Homology model building

The protocol follows a previously published method (56). Aligned protein sequences of mouse 1092 ORs are manually aligned to pre-aligned protein sequences of 11 GPCRs including bovine rhodopsin (PDB: 1U19), human chemokine receptors CXCR4 (PDB: 3ODU) and CXCR1 (PDB: 2LNL), and human adenosine a2A receptor (PDB: 2YDV) using Jalview (34).

Four experimental GPCR structures (1U19, 3ODU, 2YDV and 2LNL) are used as templates to build Olfr539 and its mutants (G154C, V209G, and L155A) and Olfr541 and its mutants (C154G and C154G/G209V) by homology modeling with Modeller. Five models are obtained and the one fulfilling several constraints (binding cavity sufficiently large, no large folded structure in extra-cellular loops, all TMs folded as α-helices, a small α-helix structure between TM3 and TM4) is kept for further molecular dynamics simulations.

### Molecular dynamics simulations

#### Olfr539/Olfr541 and their mutants systems

The models were embedded in a model membrane made up of POPC lipids solvated by TIP3P water molecules using Maestro. The total system is made up of ∼48,650 atoms in a periodic box of 91*89*98 Å^3^.

Molecular dynamics simulations are performed with sander and pmemd.cuda modules of AMBER12 with the ff03 force-field for the protein and the gaff.lipid for the membrane. Hydrogen atoms are constrained by SHAKE algorithm and long-range electrostatic interactions are handled with Particle Mesh Ewald (PME). The cutoff for non-bonded interactions is set at 8 Å. Temperature and pressure are maintained constant with a Langevin thermostat with a collision frequency of 2 ps^-1^. In addition, a weak coupling anisotropic algorithm with a relaxation time of 1 ps^-1^ is applied. Snapshots are saved every 20 ps.

Two energy minimizations are performed during 10,000 steps with the 5,000 first steps using a conjugate gradient algorithm. The first one is run with a restraint of 200 kcal.mol^-1^ applied on all atoms of the membrane and water and the second one with the same restraint on all atoms of the receptor. This last constraint is kept for the heating phase of 20 ps (NTP, 100K to 310K, Langevin thermostat with a collision frequency of 5 ps^-1^) and equilibration of 15 ns (NTP, 310K). Restraints are then reduced by 5 kcal.mol^-1^Å^-2^ and another cycle of minimization-equilibration is performed. The systems (Olfr539 models (wt, G154C, V209G and L155A) and Olfr541 models (wt, C154G and C154G/G209V)) are replicated six times and 525 ns-long production molecular dynamics are performed after an equilibration period of 50 ns. RMSDs of seven transmembrane domains were calculated using CPPTRAJ in AmberTools. The RMSDs are between initial positions and each frame in the production step. 3D structures were visualized using VMD.

#### Other Olfr pairs

We used the Membrane Builder (57) utility of CHARMM-GUI (58) for embedding each of the receptors into a pre-equilibrated simulation box of a membrane composed by POPC lipids. Each of these protein-lipid complexes was solvated in explicit TIP3P water molecules in a dodecahedron box (approximate dimension of 7.80 nm X 7.80 nm X 10.68 nm) separately and sodium and chloride counterions were added for maintaining the physiological salt concentration of each system at 150 mM. We used the software GROMACS (59) (version 2019.4) in combination with the all-atom CHARMM36 (60) force field for performing MD simulations at 310 K coupled to a temperature bath with a relaxation time of 0.1 ps (61). Pressure was calculated using molecular virial and held constantly by weak coupling to a pressure bath with a relaxation time of 0.5 ps. Each system was first subjected to a 5000 step steepest descent energy minimization for removing bad contacts (62). Then, the systems were heated for 100 ps in steps of ramping up the temperature to 310K under constant temperature-volume ensemble (NVT). Equilibrium bond length and geometry of water molecules were constrained using the SHAKE algorithm (63). We used a time step of 2 fs. The short range electrostatic and van der Waals (VDW) interactions were estimated per time step using a charge group pair list with cut-off radius of 8 Å between the centers of geometry of the charged groups. Long range VDW interactions were calculated using a cut-off of 14 Å and long-range electrostatic interactions were treated using the particle mesh Ewald (PME) method (64). Temperature was kept constant by applying the Nose-Hoover thermostat (65). Parrinello-Rahman barostat (66) with a pressure relaxation time of 2 ps was used for attaining the desired pressure for all simulations. The simulation trajectories were saved each 200 ps for analysis. The protein atoms were position restrained using a harmonic force constant of 1000 kJ mol-1 nm-2 during the NVT equilibration stage while the lipid and water molecules were allowed to repack around the protein. The system was further equilibrated at NPT by reducing the force constant on protein atoms from 5 kJ mol-1 nm-2 to zero in a stepwise manner for 3 ns each while having the pressure coupling on. We also performed an additional 10 ns of unrestrained simulation before starting the actual production run. This accounts for a total 25 ns of NPT equilibration prior to the production run. We performed three productions runs each 400 ns long starting from three independent sets of initial velocities for each system. Three independent simulations (each 400 ns long) were performed for each system.

Calculation of RMSD: Root mean square deviation (RMSD) of a particular OR from its initial structure was computed using the gmx rms utility of GROMACS. For this, only the C-alpha atoms of the seven transmembrane (TM) domains were considered and the flexible loop regions were omitted. The variance in RMSD was calculated using the block averaging method as implemented in the gmx analyze utility.

Calculation of Residual hydrophobic mismatch (RHM): For calculating the unfavorable hydrophobic interactions between protein and lipid, we took into account the TM hydrophobic residues not making sustained contacts (< 20% of the total simulation time) with either membrane tail or head groups using the module gmx select along with -om utility. The residual hydrophobic mismatch (RHM) between receptor and membrane was expressed in terms of the cumulative solvent accessible surface area (SASA) of TM hydrophobic residues (Gly, Ala, Pro, Val, Met, Cys, Ile, Trp, Phe, Tyr and Leucine) following the above criteria (67). Per residue SASA was calculated using the gmx sasa tool along with -res option as implemented in GROMACS.

### OR protein sequence analyses

Protein sequences of 1092 mouse ORs were aligned by Clustal Omega with default parameters. Conservation degree of amino acid residues was visualized by WebLogo.(68) To identify the amino acid residues involved in RTP dependence, Grantham distances (36) were calculated for all pairs of ORs at each position. Grantham distances consist in attributing distances numbers between two amino acids that are proportional to their evolutionary distance. The positions where Grantham distances of RTP-independent ORs were significantly shorter than those of all ORs were searched by one-sided t-tests followed by Bonferroni correction.

### Designing consensus ORs

Protein sequences of human ORs were downloaded from The Human Olfactory Data Explorer (HORDE) webpage (https://genome.weizmann.ac.il/horde/). The protein sequences were aligned using MAFFT. The most frequently used amino acid residues were defined as consensus residues at each position. The consensus amino acid sequences were translated into DNA sequences using Codon Optimization Tool on Integrated DNA Technologies (IDT) webpage.

### Luciferase assay in Hana3A cells

The Dual-Glo Luciferase Assay (Promega) was used to determine the activities of firefly and Renilla luciferase in Hana3A cells as previously described (56). Briefly, firefly luciferase, driven by a cAMP response element promoter (CRE-Luc; Stratagene), was used to determine OR activation levels. For each well of a 96-well plate, 5 ng SV40-RL, 10 ng CRE-Luc, 5 ng mouse RTP1s, 2.5 ng M3 receptor3, and 5 ng of Rho-tagged receptor plasmid DNA were transfected. Normalized activity for each well was further calculated as (Luc-400)/(Rluc-400) where Luc = luminescence of firefly luciferase and Rluc = Renilla luminescence. The basal activity of an OR was averaged from six wells in the absence of odorants and further corrected by subtracting that of the control empty vector. An odorant-induced activity was averaged from at least three wells and further corrected by subtracting the basal activity of that receptor. Odorant-induced responses were normalized to that of wt.

## Acknowledgments

We thank Mengjue Jessica Ni for expert technical assistance and Conan Juan, Sahar Kaleem, Aashutosh Vihani, Kevin Zhu and Yen Dinh for manuscript editing. This work was supported by grants from NIH (HM: DC014423 and DC016224, NV: NIH R01-GM097261). KI stayed at Duke University with financial support from Tokyo University of Agriculture and Technology as a student of the program for leading graduate schools in Japan. YF stayed at Duke University with financial support from JSPS Program for Advancing Strategic International Networks to Accelerate the Circulation of Talented Researchers.

## Author Contribution

HM conceived and designed the project. KI, RS, ESB, YEL and HM performed research concerning the large-scale expressions screening of OR and Olfr539 and Olfr541 mutants. KI, MD, and CADM performed the molecular modeling of Olfr539 and Olfr541 plus their mutants. MHN, YF, CADM and HM performed research on the consensus receptors. NV, SG, CADM and HM performed the salt bridge insertion. KI, CADM, MY and HM carried out the analysis and wrote the paper with inputs from all authors. HM supervised the project.

---

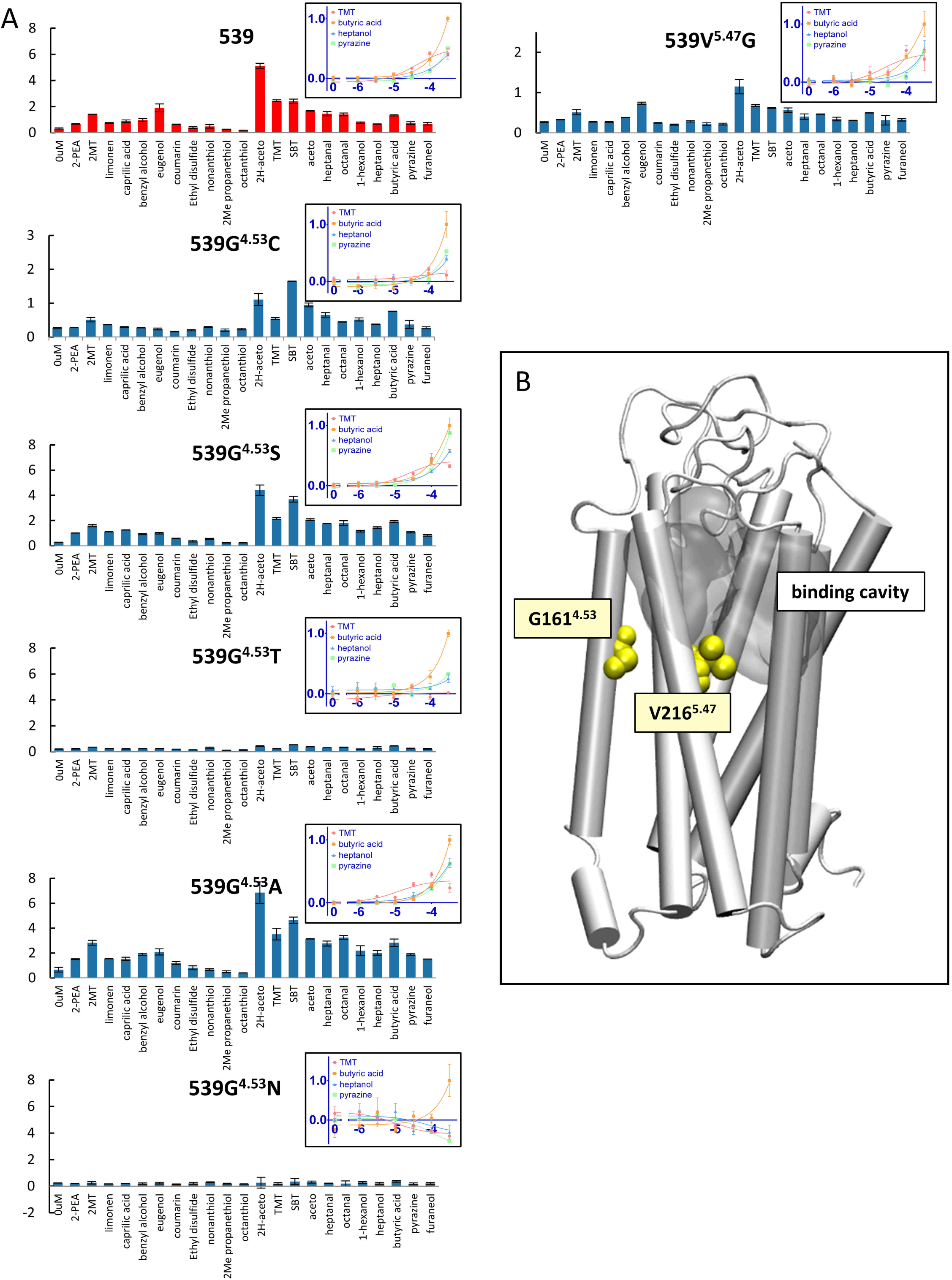

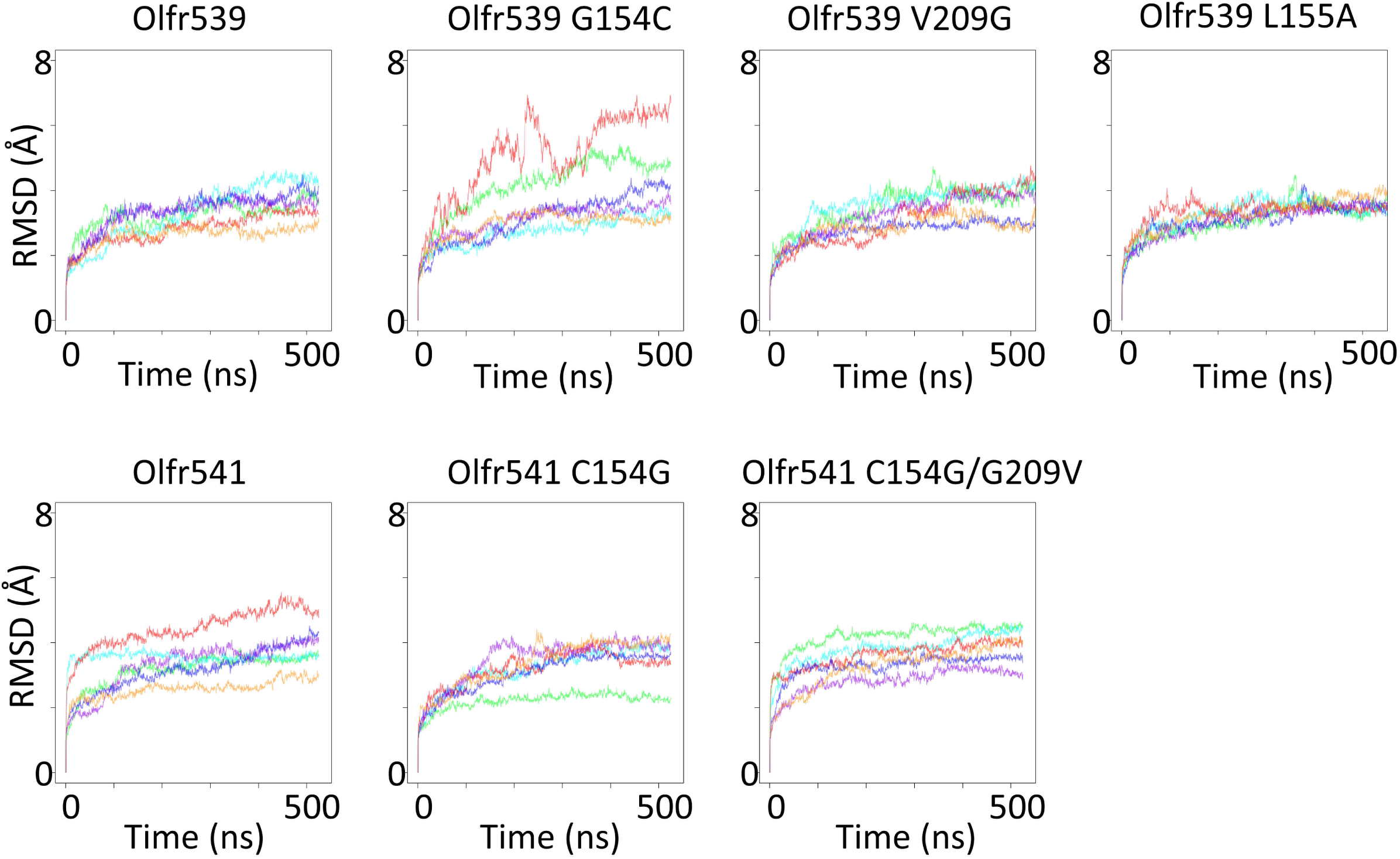

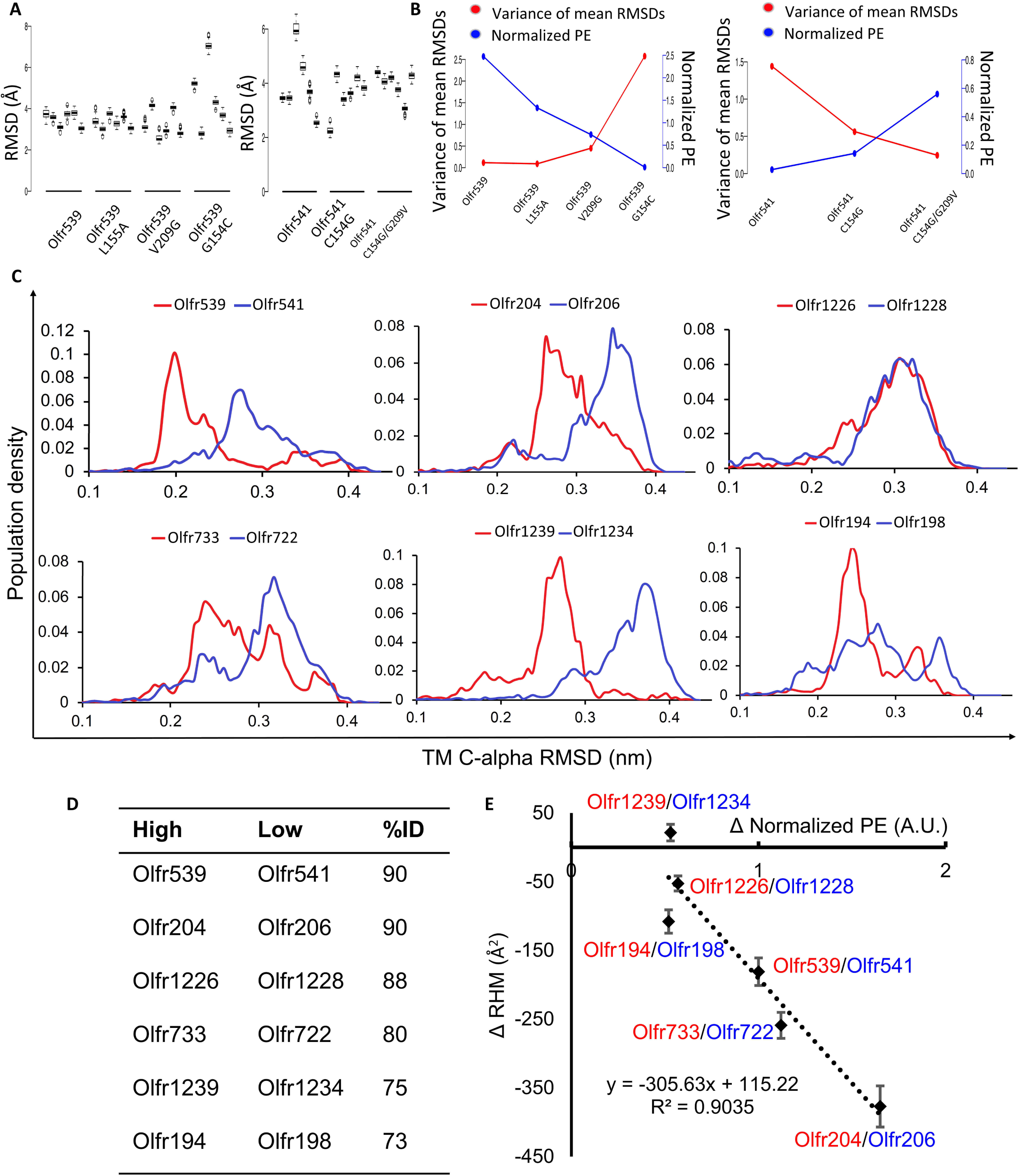

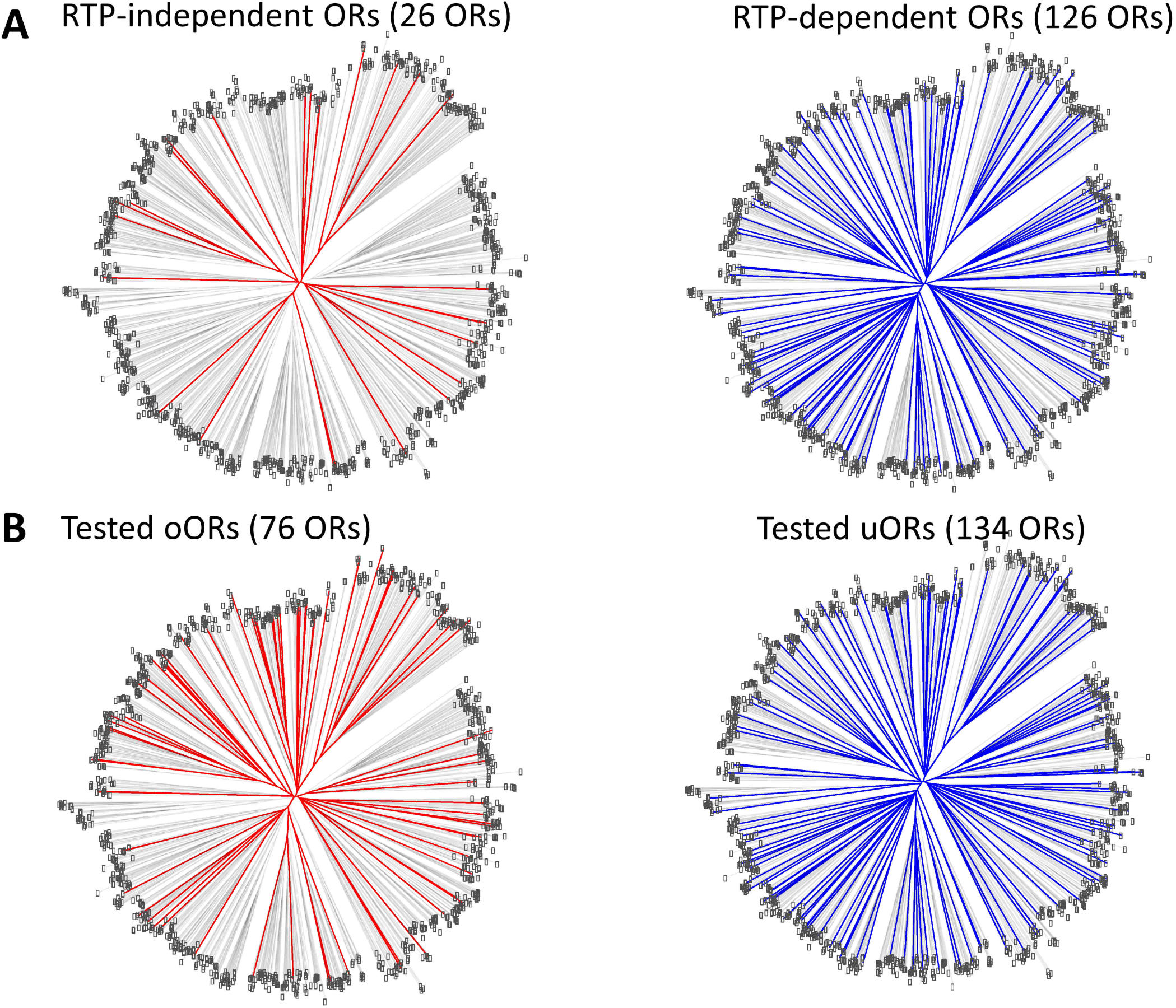

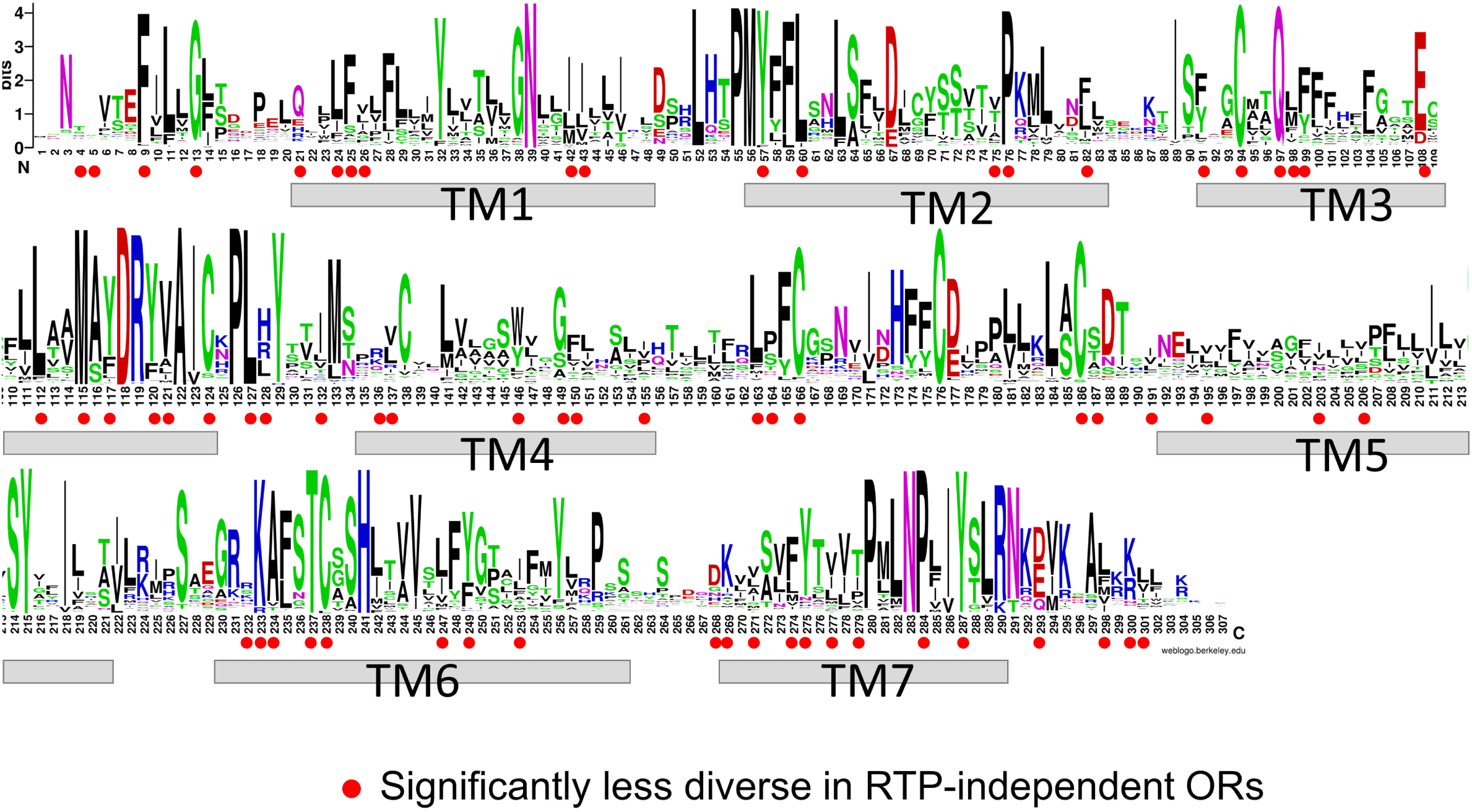

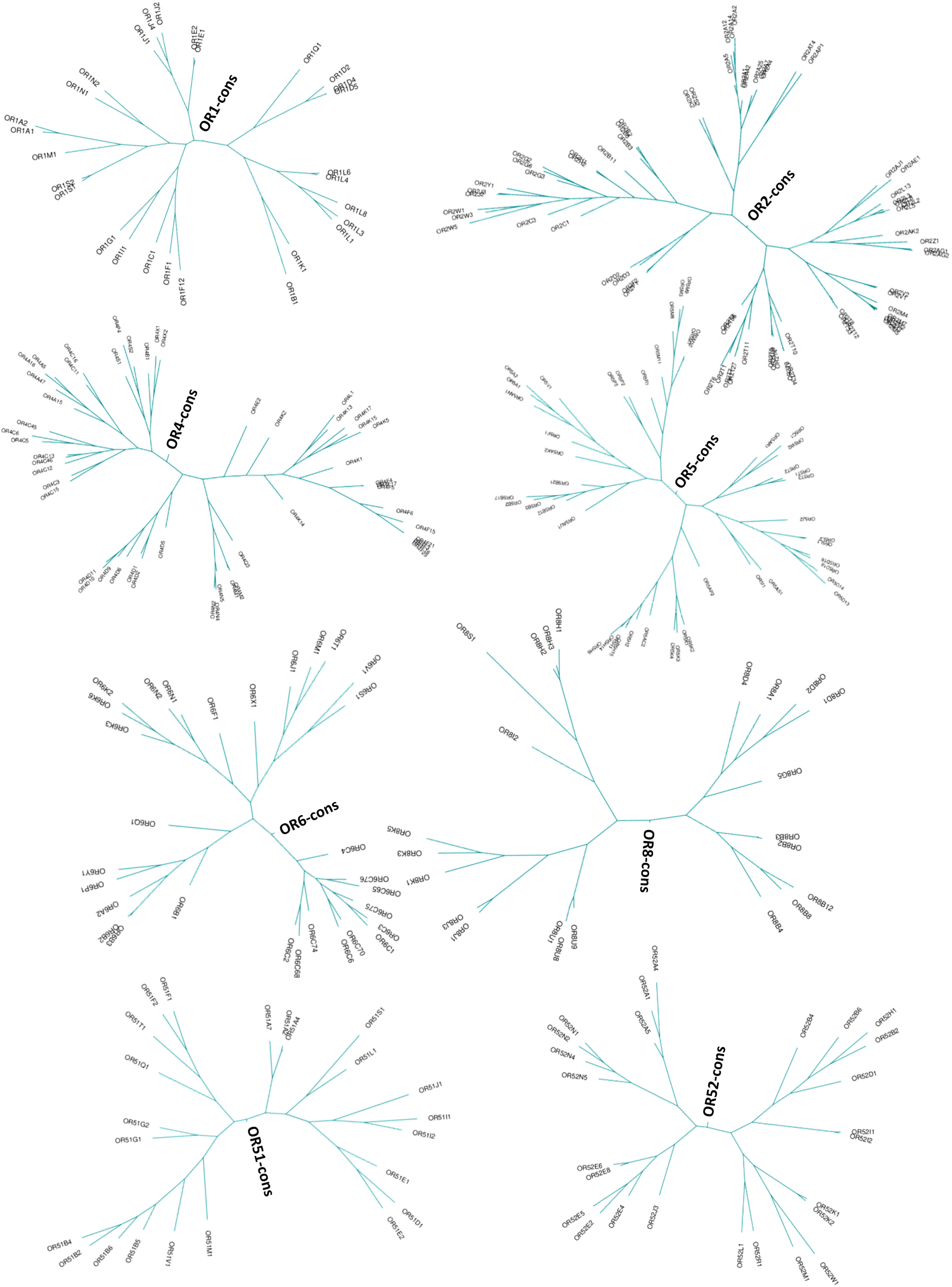

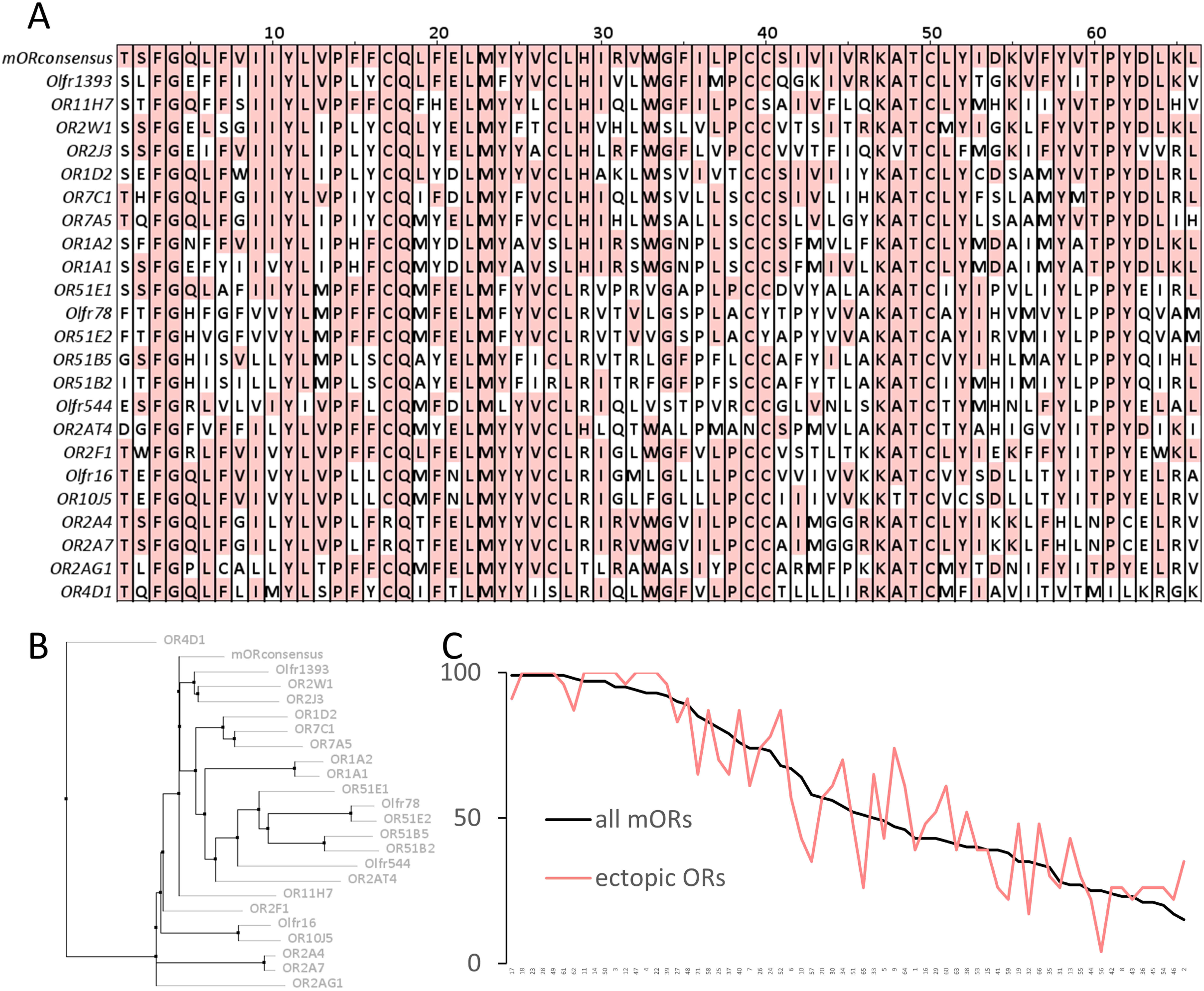

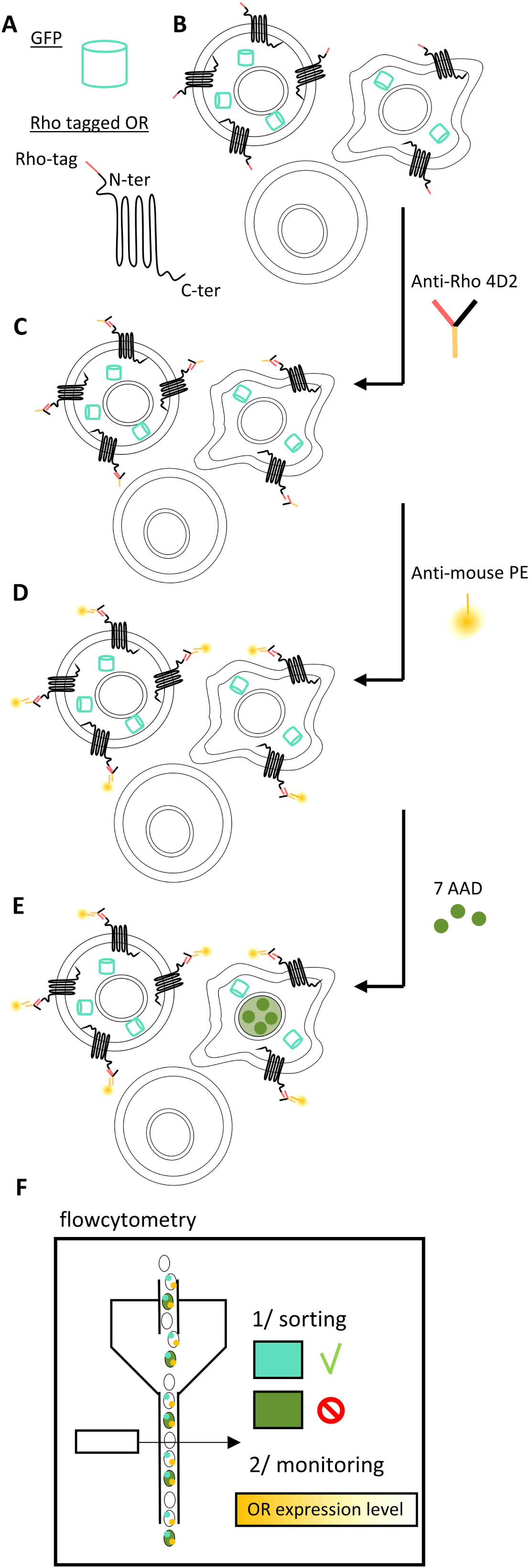

