## Supplementary information for "Structural instability and divergence from conserved residues underlie intracellular retention of mammalian odorant receptors"

**Figure S1.**

Influence of G4.53 and V5.47 on the odorant binding. **A.** Screening of the response in luciferase assay of Olfr539 and mutants (cell surface expression can be found in Fig. 1) against 22 odorants tested at 150µM. The y-axis represents the normalized luciferase luminescence and is indicative of the receptor response. Insert: dose response for each corresponding OR performed against four agonists; TMT (red), butyric acid (orange), 1-heptanol (blue) and pyrazine (green). B. Homology model of Olfr539 represented in gray cartoon. G4.53 and V5.47 are highlighted in yellow Van der Walls representation. The canonical odorant binding cavity is localized by a gray cloud.

**Figure S2.**

RMSDs of 6 individual MD simulations of Olfr539 systems (wild type, L162A^4.54^ G161C^4.53^ and V216G^5.47^) (left), and Olfr541 systems (wild type, C154G^4.53^ and C154G^4.53^/G209V^5.47^) during 600ns of MD simulation. Each simulation is represented by a line of a different color.

**Figure S3.**

**A**, RMSDs of top half (extracellular side) of transmembrane domains of 6 individual MD simulations of Olfr539 systems (wild type, L162A^4.54^ G161C^4.53^ and V216G^5.47^) (left), and Olfr541 systems (wild type, C154G^4.53^ and C154G^4.53^/G209V^5.47^) (right). The models are placed in descending order based on cell surface expression levels for Olfr539 systems and in ascending order for Olfr541 systems. **B**, Plots of the variance of mean top half RMSDs (left axis, red plots) and cell surface expression levels (right axis, blue plots) of Olfr539 systems (left) and Olfr541 systems (right). **C,** Population density distribution of RMSD of TM C-α atoms for six Olfr pairs (high expression in red and low expression in blue) over the entire 1.2 μs of protein MD trajectory. **D,** The six Olfr pairs selected that show high percentage of identity (%ID) in their amino acid sequences but different cell surface expression levels, defined here as High and Low expression. **E,** For each Olfr pair, the difference in RHM and in cell surface expression between the Olfr showing high and low cell surface expression levels were calculated (ΔRHM and ΔNormalized PE). The anti-correlation between these two parameters appears to be linear.

**Figure S4.**

**A**, Both RTP-independent ORs (left) and RTP-dependent ORs (right) are distributed on multi branches of the phylogenetic tree based on protein sequences of mouse ORs. **B**, Both tested oORs (left) and tested uORs (right) are distributed on multi branches of the phylogenetic tree based on protein sequences of mouse ORs.

**Figure S5.**

The identified 66 sites adhere to highly conserved sites. The identified 66 sites are colored (red dot) on the consensus protein sequences of mouse ORs with amino acid conservation degree.

**Figure S6.**

Ancestral tree of each human OR family including their corresponding consensus OR. Ancestral Parsimony trees are made based on the maximum parsimony score and the edge length are built according to the ACCTRAN criterion.

**Figure S7.**

Conservation of the 66 positions in ectopic human and mouse ORs. A. The 66 positions are shown for each selected ectopic ORs and the consensus sequence of mouse ORs. For each position, residues are colored in light red if they match with the amino acid of the consensus sequence. ORs are sorted following the tree presented in **B**. **B.** Tree built by % of identity of the 66 positions including the consensus mOR sequence and all selected ectopic ORs. **C.** Usage rate of consensus residues at the 66 sites for ectopic ORs (red) and all mouse ORs (black).

**Figure S8.**

Method to evaluate cell surface expression of ORs (see Method for complete description). **A.** Cells are transfected with GFP (green) and Rho-tagged (red) OR (black). **B.** Cells after transfection. They can be living transfected cells, dead partially transfected cells, or non-transfected cells. **C.** Cells after staining with an Anti-Rho 4D2 antibody. **D.** Cells after staining with an Anti-mouse PE F(ab')₂ Fragment antibody (yellow) **E.** Cells after staining with 7AAD, selective to dead cells. **F.** Flowcytometry where cells are first sorted to only select GFP positive and 7AAD negative cells. The intensity of PE fluorescence of the selected cells is then monitored to evaluate the cell surface expression of the OR.

**Table S1.**

Alignment of OR sequences used to create the consensus ORs for each subfamily.
